# Synaptic Proteomes of Cortical Interneuron Classes Revealed by Antibody Directed Proximity Labeling

**DOI:** 10.1101/2024.01.03.574066

**Authors:** Alexandria S. Battison, Jennifer C. Liddle, Stefan L. Sumsky, Christopher B. O’Connell, Jeremy L. Balsbaugh, Joseph J. LoTurco

## Abstract

Subtypes of inhibitory interneurons play diverse roles within neural circuits in cerebral cortex. Defining the molecular underpinnings of interneuron functions within cortical circuits will require identification of interneuron synaptic proteomes. In this study, we first combined genetically directed expression of tdTomato-synaptophysin with antibody-directed proximity labeling and tandem mass spectrometry to identify synaptic proteomes of three major interneuron classes in mouse cortex: parvalbumin (PV), somatostatin (SS), and vasoactive intestinal peptide (VIP).

After stringent filtering we identified 581 proteins: 228 identified in all cell classes and 353 in one or two of three classes. The PV class had the largest number of uniquely identified proteins (141), followed by VIP (30) and SST (20). Consistent with previously reported electrophysiological evidence, PV presynaptic proteomes were enriched for NMDA receptor subunits and scaffolding proteins. We used antibodies against synaptotagmin 2 (Syt2), a presynaptic protein present at PV synapses, to confirm NMDAR localization, and to find that the mu-opioid receptor agonist buprenorphine rapidly caused reorganization of the PV presynaptic proteome. Overall, our results reveal proteomes of PV, SST, and VIP interneurons in cortex that likely underlie distinct and dynamic interneuron synaptic properties.

## Introduction

The brain is comprised of diverse neuron types defined by morphology, connectivity, physiology, transcriptomes, and proteomes. Different neuron types form synapses that can be distinct in physiology, neurochemistry, and morphology, and this diversity is determined and maintained by the complement of proteins present at synapses. The vertebrate neocortex is predominantly comprised of excitatory glutamatergic neurons and inhibitory GABAergic interneurons. Inhibitory interneurons in the cerebral cortex are exceedingly diverse, with over 20 distinct subtypes defined ^1–3,47^. GABAergic interneurons provide a break on excitation, control timing and synchrony within neural circuits^4–7^, and can be modulated differentially according to cell class^8,9^. Moreover, disruptions in GABAergic interneurons in cortex have been implicated in several neurological conditions including Alzheimer’s Disease^10–12^, schizophrenia^13,14^, autism spectrum disorder^15,16^, and epilepsy^17–19^.

Three major classes of interneurons in cortex are partially defined by their expression of parvalbumin (PV), somatostatin (SST), or vasoactive intestinal peptide (VIP)^1,20,21^.These three classes are distinct in their morphology and connectivity, and play specialized roles in neocortical circuits^8^. PV interneurons perform fast inhibition and project to the soma and proximal dendrites of excitatory pyramidal cells (PCs)^7,22^. This perisomatic innervation provides tight control of PC spike timing and can quickly alter PC cell firing, thus dampening excitation within circuits^23^. Somatostatin interneurons, in contrast, project to distal dendrites and predominantly facilitate and perform uniform suppression of PC spiking^6^. In contrast to the other two classes, VIP interneurons synapse predominantly onto other inhibitory cells^24^. In doing so, VIP cells are disinhibitory and indirectly potentiate excitatory circuits^24,25^.

Extensive characterization of transcriptomic profiles of inhibitory GABAergic interneurons in rodent cerebral cortex has revealed multiple differences between interneuron cell classes at the molecular level^3^. While such transcriptomic profiling has provided critical information regarding the molecular identity of these interneurons, RNA expression does not provide direct information on protein expression, protein localization, or protein-protein interactions. As protein-based mechanisms are the primary functional units within cells, proteomic characterization is essential to reveal molecular pathways operating within different interneuron synapse types.

Obtaining cell-type and cell-compartment defined proteomes by classical biochemical approaches in the central nervous system has previously posed challenging. Cell-types can be specifically labeled in mouse models by genetically directed expression of Cre recombinases and/or fluorescent reporters, and labeled cells can then be isolated by fluorescent cell sorting. However, cell dissociation methods result in the loss of many subcellular compartments including presynaptic terminals. Subcellular compartments such as synapses can be isolated through centrifugation and fractionation; however, information on cell type is typically lost after subcellular fractionation. Proximity labeling methods, including BioID, TurboID, and APEX^26–28^, have emerged as powerful approaches to identify subcellularly localized proteins at synapses and other cellular compartments of defined cell types^10,29,30^. In proximity labeling, biotinylation occurs through the covalent addition of biotin to exposed lysine (BioID) or tyrosine (APEX) residues. The result is a radius of biotinylated proteins in proximity to bait proteins, and the biotinylated proteins can then be enriched by binding to streptavidin and identified via liquid chromatography tandem mass spectrometry. Proximity labeling methods have been used to identify subcellular proteomes and more recent advances in proximity labeling have shown great success in obtaining cell-type specific neuroproteomes^10^.^31^ For example, APEX2 proximity labeling has successfully been used to identify cell-type and cell-compartment specific local proteomes of striatal neurons^10^. In addition to genetically encoded methods such as APEX and BioID/TurboID, antibody-based proximity labeling methods have also been developed, in which bait proteins are labeled with primary antibodies and then biotinylation of nearby proteins is achieved through peroxidase activity of HRP conjugated secondary antibodies^31,32^.

In this study, we used a modified antibody-based proximity labeling method to obtain interneuron class and presynapse-specific proteomes from three major subtypes of GABAergic cortical interneurons. We crossed mouse lines that express Cre-recombinase in PV, SST, and VIP interneurons ^33,34^ with a line that conditionally expresses a synaptically localized fusion protein, tdTomato:Synaptophysin. Antibodies against tdTomato were then used to direct proximity labeling in interneuron classes. The proteomes obtained with this method displayed both overlapping and unique features. Shared features included presynaptic vesicle proteins and proteins known to mediate exo- and endocytosis. In addition to shared pathways, the PV presynaptic proteome was uniquely and significantly enriched for NMDA receptor subunits and NMDAR scaffolding proteins, and this was confirmed with a second antibody against the PV presynapse-associated protein synaptotagmin 2 (Syt2). We used stimulated emission depletion (STED) super-resolution microscopy to confirm that presynaptic NMDA receptors are redistributed in response to the administration of the mu-opioid partial agonist, buprenorphine. Together our study used a novel antibody based proximity labeling approach to characterize synaptic proteomes of interneurons, and demonstrated that antibody-based proximity labeling can capture dynamic changes to interneuron-class specific proteomes.

## Results

### Antibody-based proximity labeling identifies VIP, SST and PV interneuron proteomes

In order to obtain cell-class specific synaptic proteomes, we generated three lines of mice expressing a tdTomato-synaptophysin fusion protein in PV, SST or VIP interneurons (Fig 1A, B). Following perfusion, fixation, cortical dissection, and vibratome sectioning (60 μm) we performed antibody-based proximity labeling on 6 brains from each mouse line (N=18 total, 6 per cell-class) (Fig 1C). Antibody-based proximity labeling by biotinylation was achieved by sequentially adding a primary antibody directed against the cell-type specific tdTomato:Synaptophysin antigen, an HRP-conjugated secondary antibody, followed by incubation with a lab-synthesized biotin-tyramide and hydrogen peroxide to catalyze biotinylation of tyrosine residues proximal to the tdTomato:Synaptophysin bait protein (Fig 1D). Successful biotinylation was validated by immunohistochemistry in all three interneuron cell-classes by the presence of FITC-streptavidin staining colocalized with tdTomato signal (Fig 1E).

**Figure 1:**
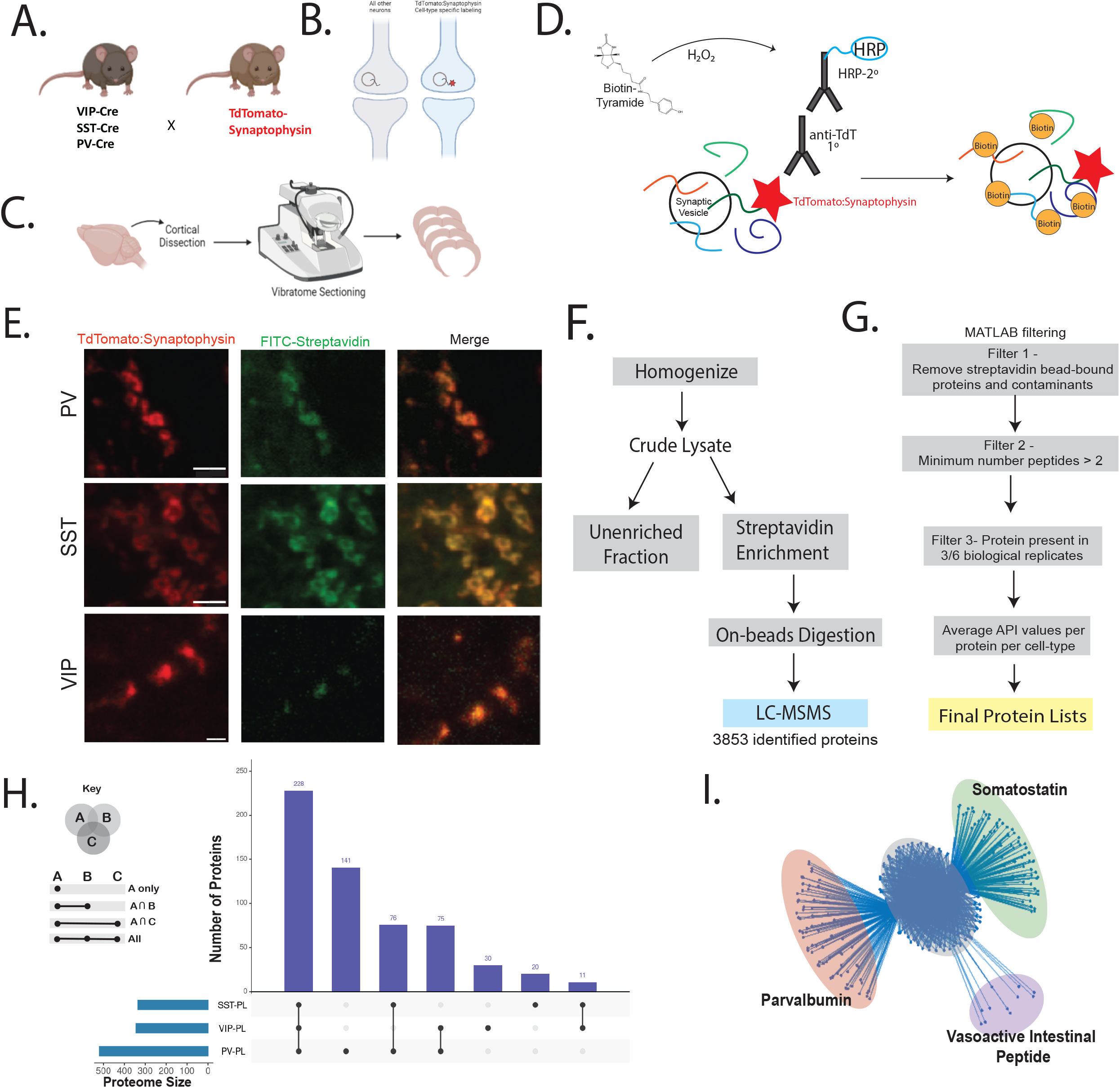
Interneuron class specific proteomes isolated by genetically directed proximity labeling in mouse cortex A) GABAergic cell-type specific cre recombinase expressing mouse lines were crossed with a tdTomato:synaptophysin fluorescent reporter mouse line to generate the three lines used in this study. B) Depiction of the resulting expression of tdTomato-synaptophysin in the synaptic terminals of select interneurons. C) Schematic of workflow for preparation of paraformaldehyde fixed, microdissected, and sectioned cortical tissue for use in proximity labelling. D) Conceptualization of antibody-based proximity labeling resulting in biotinylation of proteins in proximity to tdTomato-synaptophysin. E) Laser scanning confocal images of tdTomato and FITC-streptavidin showing the colocalization of biotinylated proteins and the bait protein tdTomato-synaptophysin (scale bar = 10um). F) Workflow for enrichment of biotinylated proteins and bottom-up label free mass spectrometry. G) Peptide search and protein processing and filtering workflow included the removal of contaminants and bead-bound endogenously biotinylated proteins, as well as computational filtering. H) UpSet plot (https://intervene.shinyapps.io/) displaying the numbers of shared and unique proteins identified after filtering for PV-PL, SST-PL, and VIP-PL datasets. I) Graph theory condensation of an unbiased, undirected network resulting in a de-novo grouping of the three inhibitory cell classes from PV-PL, SST-PL and VIP-PL proteomes.

Following successful biotinylation, tissue sections were homogenized, and biotinylated proteins were enriched by streptavidin pull-down (one brain /pull down), with no pooling of samples. Proteins were then analyzed by liquid chromatography tandem mass spectrometry (LC-MSMS). To prepare samples for bottom-up shotgun proteomics, samples were reduced and alkylated, and biotinylated peptides were digested off beads with trypsin (1:20, Promega) (Fig 1F). Peptide sequences were identified based on MS2 spectra acquired in the linear ion trap, and raw data were processed using MaxQuant v1.6.0.1 and search results were filtered to a 1% false discovery rate (FDR). A total of 3853 proteins were identified across all 18 samples prior to additional filtering (Supplemental Data Table 1). The dataset was then further filtered to remove proteins identified by only 1 peptide (2258 proteins removed), and any proteins that were present in fewer than 3/6 replicates in any cell type (1014 proteins removed). The final list contains 581 high-confidence, reproducibly observed proteins (Fig 1G). 141 proteins were uniquely identified by proximity labeling in parvalbumin presynaptic terminals (PV-PL samples), 20 in somatostatin presynaptic terminals (SST-PL samples), and 30 in vasoactive intestinal peptide proximity labeled presynaptic terminals (VIP-PL samples). Seventy-five proteins were shared between PV and VIP, 76 between PV and SST, 11 between SST and VIP, and 228 were present in all three cell types. An UpSet plot (Fig 1H) displays the intersection of proximity labeled proteins across all three classes of cells (PV-PL, SST-PL, and VIP-PL). In order to determine whether the proteins identified in the three proximity labeling conditions clustered according to three cell classes, we performed unbiased and unsupervised network analysis of shared proteins. We found that the network analysis resulted in 4 clusters of proteins, one with proteins shared by all conditions and three additional clusters formed by proteins elevated in each cell class respectively (Fig 1I). This unbiased clustering approach suggests that we are able to successfully capture and preserve proteome level differences between our initial three interneuron cell classes.

### Enrichment of cell compartments and pathways in PV, SST, and VIP proteomes

To test whether our approach enriched for proteins present at synapses, we performed Gene Ontology analysis (Panther) on PV-PL, SST-PL, and VIP-PL proteomes. We found that proteins in each of the three proximity labeling conditions were significantly enriched for the GO:CC (cellular compartment) terms presynapse, synaptic vesicle membrane, and presynaptic active zone compared to crude lysate, which enriched for the non-synaptic term neuronal cell body (Fig 2A). Similarly, PV-PL, SST-PL, and VIP-PL proteomes were enriched for the GO:BP (biological process) terms synapse organization, regulation of neuronal synapse plasticity, exocytosis, and synaptic vesicle priming, thus confirming that our enrichment was for the presynaptic terminal of the neuron (Fig 2B). Examining each cell-class proteomes gene ontology networks in more detail with Cytoscape,^35^ we found cellular compartment gene ontology terms for presynapse, axon, and synaptic vesicle as terms shared across all 3 local proteomes (Fig 2C-E). Cytoscape analysis also revealed cellular compartment gene ontology terms that were uniquely enriched in one cell-class, such as an enrichment for cellular-membrane proteins in the SST-PL proteome(Fig 2D). Similarly, a unique enrichment for postsynaptic density and NMDA receptors was found in PV-PL proteomes (Fig 2C). KEGG molecular signaling pathways for long-term potentiation and depression, as well as several terms related to addiction, appeared in PV-PL proteomes (Fig 2F) (Supplemental Data Table 2). In contrast, SST-PL terms included several molecular signaling pathways including PI3K/Akt/mTOR and Erbb2/Ras/MAPK (Fig 2G). The VIP-PL proteins were in terms mostly shared by the other two proteomes but also contained proteins in the gastric acid secretion pathway (Fig 2H).

**Figure 2:**
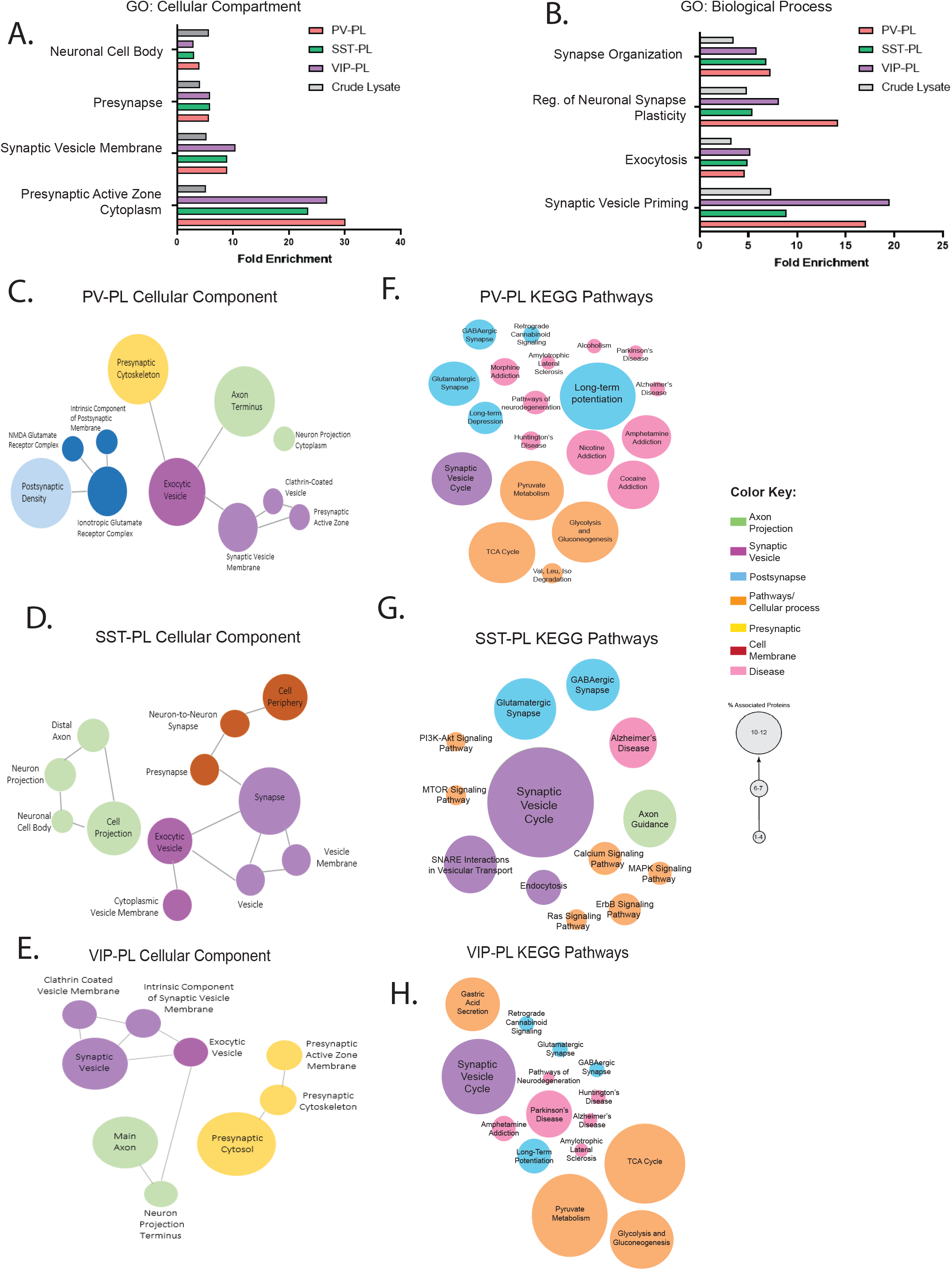
Gene ontology analysis of PV-PL, SST-PL and VIP-PL proteomes indicates enrichment for presynaptic compartments and pathways A) Panther cellular component gene ontology (GO:CC) analysis for proteins in PV-PL, SST-PL, and VIP-PL proteomes relative to crude lysate. B) Enrichment of Panther Biological Process gene ontology (GO:BP) terms for the three proximity labelled proteomes relative to non-enriched crude lysate. C) Cellular Component gene ontology networks for PV-PL. D) SST-PL and E) VIP-PL presynaptic proteomes obtained from Cytoscape network analysis with minimum number of genes/term = 2 and displaying enriched terms p< 0.05 relative to the mouse genome. KEGG molecular pathways enriched in F). PV-PL G). SST-PL H). VIP-PL. Color key and circle size legend for C-H also shown.

### PV-PL proteomes are enriched for NMDA receptors and NMDA receptor interacting proteins

Our Gene Ontology results above showed surprising enrichment in PV-PL for several terms related to the post-synapse and more specifically to glutamate receptor signaling. In order to obtain a more detailed assessment of this enrichment in glutamatergic postsynaptic mechanisms, we turned to the expert-curated synapse protein localization tool SynGO (syngoportal.org). While all three interneuron classes had proteins present from both pre- and postsynaptic compartments (Fig 3A-C), PV-PL and SST-PL samples showed a significantly higher ratio (Fig 3D) of post-synaptic proteins relative to presynaptic proteins compared to VIP-PL [ANOVA; F(2,15)=5.77, p=0.0138; Tukey PV-PL vs SST-PL, p=0.023, SST-PL vs VIP-PL, p=0.0228]. As there were over 40 proteins in PV-PL contained within postsynaptic densities (Fig 3A), we examined these in more detail (Fig 3E). PV-PL samples included several proteins associated with NMDAR signaling: Cnksr2, Dlg2, Grin2B, Iqsec2, Lrrc4b, Lrrc7,and Rtn4 (Fig 3E). Baiap2 and Dlg2 were identified in all three cell classes; however, co-occurrence was most frequent in PV-PL cells (Fig 3F, G). As Grin2B and Grin2A were unique to PV-PL, we compared the spectral counts of Grin1, Grin2A, and Grin2B across cell classes for each biological replicate. We found that total Grin1 spectra identified was greatest for PV-PL, identified in 5 out of 6 replicates (Fig 3H). Furthermore, Grin2A spectral counts were only identified in PV-PL samples, and only a single biological replicate of SST-PL contained Grin2B spectra (Fig 3I,J). Together these patterns indicate that NMDA receptor subunits and associated proteins are enriched in the PV-PL samples.

**Figure 3:**
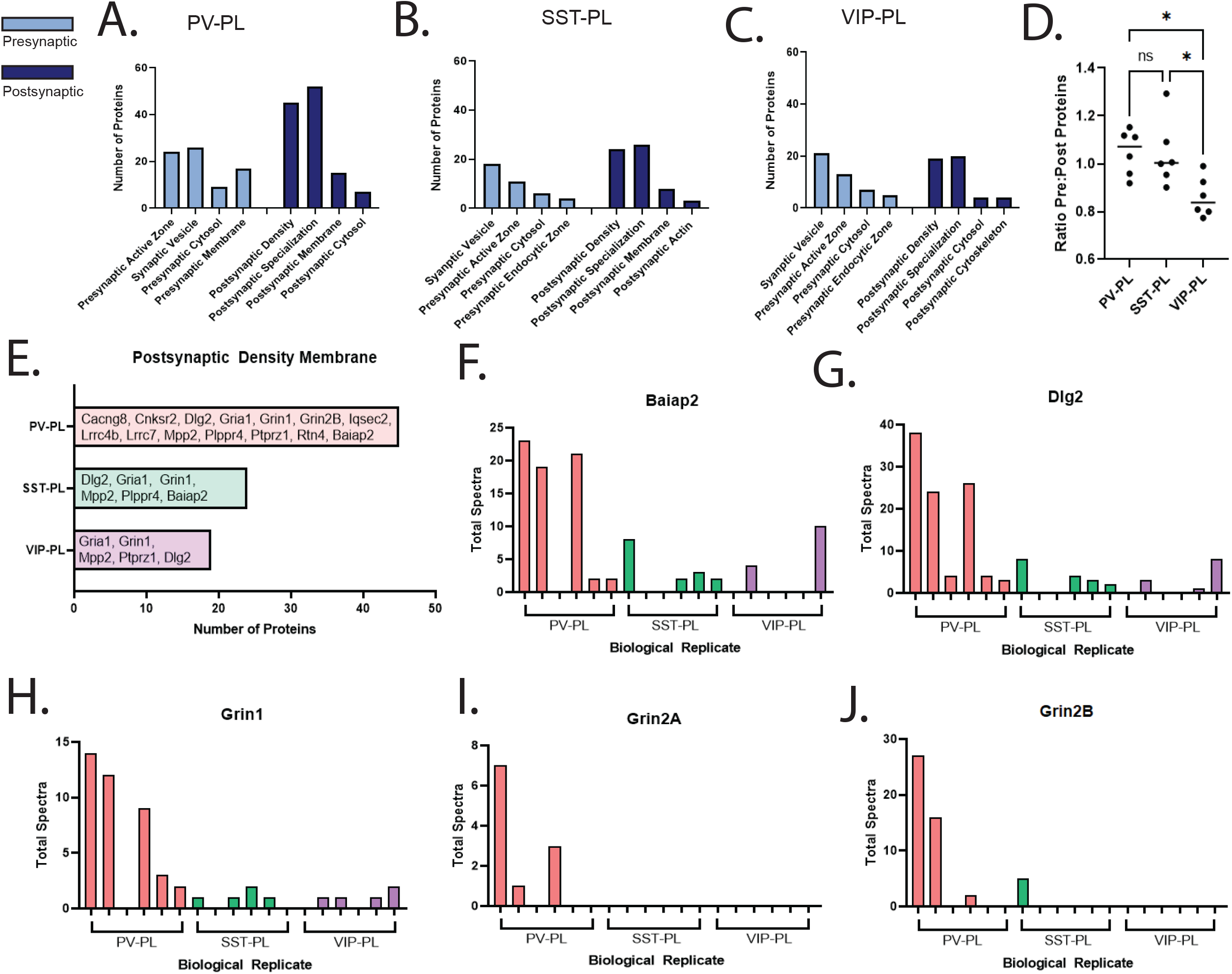
Enrichment for postsynaptic proteins and NMDARs in PV-PL proteomes. A-C) Bar graphs of the number of pre- and postsynaptic proteins in 4 SYNGO categories (Syngoportal.org) for A). PV-PL, B). SST-PL, and C). VIP-PL proteomes. D) Plot of the ratios of proteins localized to presynaptic and postsynaptic sites in each proximity labelling condition [ANOVA; F(2,15)=5.77, p=0.0138; Tukey, PV-PL vs SST-PL, *p=0.023, SST-PL vs VIP-PL, *p=0.0228]. E) Bar graph of the total number of postsynaptic proteins and NMDA receptor related postsynaptic proteins identified in PV-PL, SST-PL, and VIP-PL proteomes. F-J) MS2 spectra totals identified for Baiap2 (F), Dlg2 (G), Grin1 (H), Grin2A (I) and Grin2B (I) across all biological replicates.

### Synaptotagmin-2 colocalizes with presynaptic NMDA receptors

In order to further assess the potential localization of NMDARs to PV presynaptic terminals we used a synaptotagmin-2 (Syt2) primary antibody for both proximity labeling and STED super-resolution microscopy. Syt2 is a member of the synaptotagmin family of synaptic proteins which facilitates calcium-dependent fusion of synaptic vesicles and presynaptic membranes^37^ (Fig 4A), and it has been shown to localize to PV synaptic terminals in cortex and hippocampus^38^. We show by STED superresolution microscopy that Syt2 is present in PV-Cre:tdTomato-synaptophysin terminals (Fig 4B). Proximity labeling experiments using a Syt2 primary antibody in sections from three mice (C57BL/6) (Fig 4C) further showed that the resulting SYT2-PL proteome is similar to the PV-PL proteome (Fig 4D). As with PV-PL proteomes, SYT-PL proteomes are significantly enriched for the GO:CC terms presynaptic active zone, exocytic vesicle membrane, synaptic vesicle membrane, and post-synaptic specialization (Fig 4E). Furthermore, we investigated the localization of Grin1 relative to Syt2 by STED. Multiple terminals show signal for both Syt2 and Grin1, consistent with our observation of Grin1’s presence in the PV-PL pre-synaptic proteome (Fig 4F). The fluorescent signal of Grin1 and Syt2 within terminals was more completely overlapping (Fig 4F) than the pattern observed for Syt2 and tdTomato-synaptophysin within terminals (Fig 4B), suggesting that Grin1 may be in close proximity to the point of synaptic vesicle and presynaptic membrane fusion in PV cell synapses.

**Figure 4:**
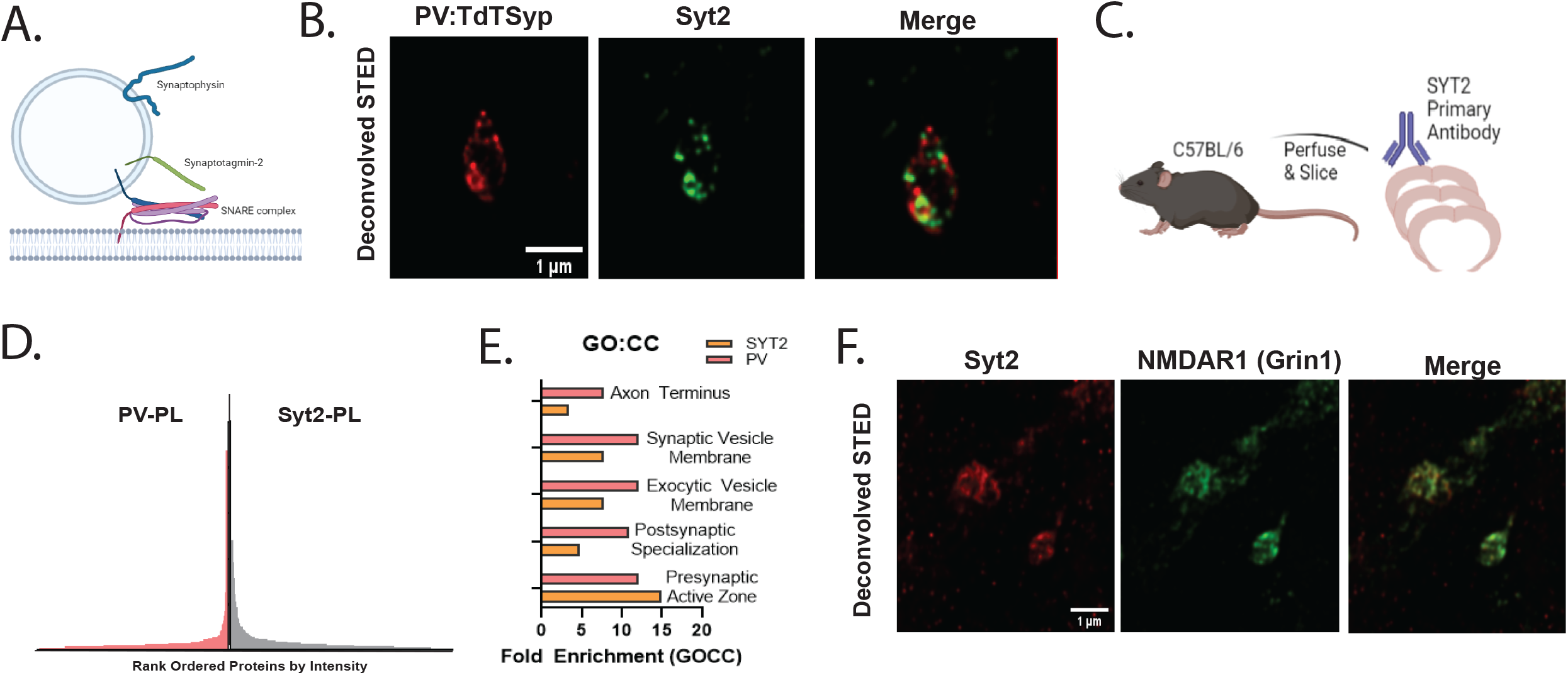
Proximity labelling and STED with Syt2 antibodies indicates NMDA receptors in PV terminals. A). Schematic of synaptotagmin-2 and synaptophysin on the synaptic vesicle (Schematic made using BioRender.com). B) STED superresolution microscopy of tdTomato (red) and Syt2 immunoreactivity (green) in PV-cre/tdTomato-synaptophysin (PV:TdTSyp) mouse cortex. C). Schematic of Syt2 antibody proximity labeling performed on C57Bl/6 mice. A total of 6 biological replicates were used. D) Mirror plot comparing rank-ordered intensities of PV-PL and SYT2-PL proteomes. E) PV-PL and SYT2-PL proteomes enrich for similar Panther Gene Ontology Cellular Components (GO:CC). I) STED microscopy images of Syt2 (red) and NMDA/Grin1 (green) show colocalization of NMDA receptors and Syt2. Scale bars in B and F = 1micron.

### Buprenorphine alters NMDAR and SYT2 colocalization and molecular proximity

Our gene ontology analysis of PV-PL proteomes indicated enrichment for several pathways related to addiction (Fig 2F; nicotine, amphetamine, and cocaine), and we performed an additional gene ontology analysis using Qiagen Ingenuity Pathway Analysis (IPA^tm^) to investigate whether additional potentially enriched pharmacological pathways are present at PV terminals. In addition to synaptogenesis and calcium signaling pathways, IPA analysis revealed that SYT2-PL proteomes contained proteins implicated in opioid signaling pathways (Fig 5A). Presynaptic effects of mu-opioid receptor activation on PV interneuron transmitter release have been shown previously^39^, and we hypothesized that mu-opioid receptor activation by the partial agonist buprenorphine could alter SYT2-PL proteomes. Our rationale was that pharmacological activation of presynaptic opioid receptors on PV cells would activate a signaling cascade that could alter presynaptic protein localization which would be detectable by both proximity labeling and STED. We administered 0.05 mg/ml of the mu-opioid high-affinity partial agonist buprenorphine or saline to C57BL/6 mice at postnatal day 30 (Fig 5B, n=3 per condition). Biotinylation in molecular proximity to Syt2 was confirmed by double labeling for Syt2 and FITC-Streptavidin (Fig 5C). Proteomic analysis of Syt2-PL revealed after filtering a total of 70 proteins unique in buprenorphine-treated mouse cortices, 167 proteins in cortices from saline-injected controls, and 128 shard between them. (Fig 5D) (Supplemental Table 4). SYNGO analysis for the synaptic compartments of proteins identified in the two conditions indicated a decrease in postsynaptic proteins in the buprenorphine condition (Fig 5E). Consistent with this analysis, LC/MS data also showed a decrease in the average precursor intensities for Grin1 and Dlg1 in the buprenorphine-treated samples (Fig 5F). Using STED microscopy and double labeling for Syt2 and NMDAR1 we found colocalization of both fluorescent signals in the presynaptic terminals of saline-treated mice (Fig 5G). Fluorescence intensity versus position plots indicate that SYT2 and NMDAR1 intensity peaks are in overlapping positions in X-Y and X-Z projections in saline-treated controls (Fig 5 H,I upper panels). In contrast, in buprenorphine-treated samples, fluorescence intensity peaks are shifted relative to each other in both projections (Fig 5 H,I lower panels). The STED microscopy results support the proximity labeling results and indicate that mu-opioid receptor activation by buprenorphine alters the proximity of SYT2 and NMDAR1 in PV synaptic terminals.

**Figure 5:**
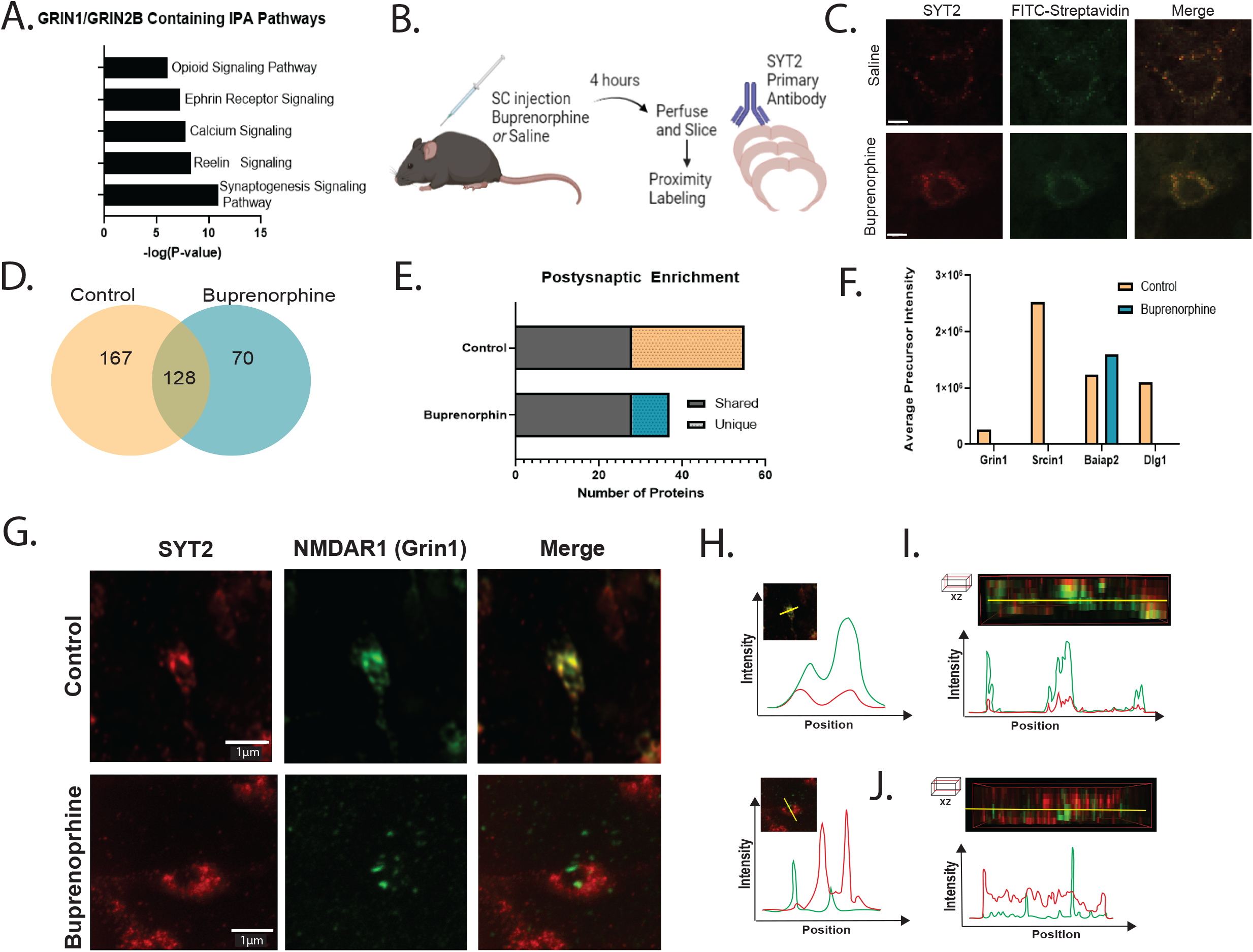
Buprenorphine shifts NMDA receptor and SYT2 colocalization and molecular proximity. A) Ingenuity Pathway Analysis (IPA) of PV-PL. IPA pathways which contain Grin1 and are enriched in PV-PL proteomes at a significance threshold of p< 0.001. B) Schematic of buprenorphine and saline experiment in which SYT-PL proteomes were captured 4 hours after drug administration. Immunohistochemistry validation of proximity labeling. C) Standard confocal microscopy image of colocalized Syt-2 immunoreactivity (red), and biotinylated proteins labeled by FITC-streptavidin in sections from both saline (upper) and buprenorphine (lower) injected conditions. D) Venn-diagram showing the total number of proteins identified after filtering in buprenorphine and saline injected animals following filtering. E) Number of unique and shared post-synaptic proteins from SYNGO in each condition (n=3). Note the loss of postsynaptic proteins from buprenorphine SYT-PL proteomes relative to saline injected controls. F) Average Precursor Intensity values of postsynaptic proteins from (E) shows a decrease caused by buprenorphine. G) 2-channel STED microscopy of Syt2 (red) and NMDAR1 (green) in buprenorphine and control conditions. Merged channels (right) show colocalization in the saline treated sample and decorrelation of the two fluorescent signals in the buprenorphine conditions. Scale bar = 1 micron. H) Fluorescence intensity vs. position graphs of XY and Z projections. Note the greater colocalization of Syt2 and NMDAR1 in control (top) compared to buprenorphine (bottom). I) Fluorescence intensity vs position for XZ planes for control and J). Buprenorphine samples.

## Discussion

In this study, we used antibody-based proximity labeling to identify presynaptic proteomes from three major classes of GABAergic cortical interneurons. The proteomes we identified in proximity to synaptophysin are significantly different between the three classes and are consistent with the unique connectivity and function of PV, SST, and VIP classes. Our dataset serves as the first interneuron cell class-specific presynaptic proteome for the cortex. By gene ontology and IPA analysis, we found that PV, SST, and VIP interneurons in the cortex have a shared ensemble of proteins involved in maintaining synaptic transmission and synaptic vesicle cycling. We also found class-specific proteins and pathways. We identified a strong presynaptic NMDAR signaling signature at the synapses of PV cells by both synaptophysin and synaptotagmin2 proximity labeling. Finally, we show that synaptotagmin-directed proximity labeling identifies a pharmacologically induced change in synaptic protein distribution following buprenorphine administration. Our strategy for antibody-based proximity labeling supports cell-type and cell-compartment specific labeling and can be leveraged to detect functional changes in protein expression following pharmacological manipulations. While we demonstrate the utility of this method in cortical synapses, this method is modular and not dependent on genetically encoded substrates, so it can be applied to other cellular compartments in diverse tissues and organs.

We observed enrichment of postsynaptic proteins in PV presynaptic terminals relative to SST and VIP terminals. This enrichment included NMDA receptor subunits Grin1, Grin2A, and Grin2B. Consistent with these findings, previous electrophysiology and electron microscopy experiments have found evidence for presynaptic NMDA receptors at PV presynaptic terminals ^40,41^. Presynaptic NMDA receptors have also been implicated in spike timing^42^ plasticity and activity-dependent plasticity ^43,44^. Interestingly, we also identified Rimbp in PV terminals, a protein shown to act downstream of calcium^45^ to facilitate vesicle association with the active zone in cerebellar interneuron synapses. Similarly, we identified Src-kinase^46^ and several proteins in PV presynaptic terminals involved in synaptic plasticity. Consistent with a possible role for changes in localization of preNMDARs in modulation or plasticity at PV terminals, we found that NMDARs at PV termini change in localization after mu-opiod receptor activation. A reduced association of presynaptic NMDARs with the active zone would be expected to alter calcium influx and change the release of transmitter. Such changes in pre-synaptic NMDAR localization may be an underappreciated mechanism of synaptic modulation and plasticity.

Mu-opioid receptors are located throughout the brain, including on the presynaptic terminal of PV interneurons. Buprenorphine is a known mu-opioid partial agonist and clinically is administered in combination with naloxone for opioid-abuse disorders. Following administration of buprenorphine and subsequent proximity labeling of PV terminals using the SYT2 primary antibody, we noticed a stark decrease in postsynaptic enrichment that appeared to be driven in part by a loss of the NMDA receptor subunits Grin1, Grin2A, and Grin2B. Based on the change in position observed by STED microscopy, we infer that this decrease in LC-MS/MS detection of NMDA subunits following buprenorphine is due to altered localization of NMDA that results in it no longer being within the radius of biotinylation. Along with a loss of NMDAR subunits at PV synapses, we also note an overall decrease in molecular pathways associated with synaptic function and neurotransmission after buprenorphine. This suggests that presynaptic modulation of PV terminals by mu-opioid receptors occurs by the redistribution of NMDA receptors.

## Supporting information

Supp. Table 1: Cell Type MSMS Data

Supp. Table 2: KEGG

Supp. Table 3: IPA Pathways

Supp Table 4: Buprenorphine

## Acknowledgements

The authors would like to acknowledge the NIH S10 High-End Instrumentation Award 1S10-OD028445-01A1, which supported this work by providing funds to acquire the Orbitrap Eclipse Tribrid mass spectrometer housed in the University of Connecticut Proteomics & Metabolomics Facility. Additionally the authors would like to acknowledge the NIH grant S10OD023618 awarded to Christopher O’Connell for support to provide funds to acquire the Abberior Instruments Expert Line STED Superresolution Microscope.

## Online Methods

### Transgenic mouse lines

All mice were housed in compliance with the Institute for Animal Care and Use Committee at the University of Connecticut. In order to target specific GABAergic interneuron cell types, VIP-IRES-Cre (VIP^tm1^(cre)^Zjh^/J JAX stock no 010908), SST-IRES-Cre (Sst^tm2.1^(cre)^Zjh^/J, JAX stock no 013044) and PV-Cre (Pvalb^tm1^(cre)^Arbr^/J, JAX stock no 017320) were used. VIP-IRES-Cre, SST-IRES-cre, and PV-Cre mice were crossed to a tdTomato-Synaptophysin (Gt(ROSA)26Sor^tm34.1^(CAG–Syp/tdTomato)^Hzs^/J, JAX stock no 012570) reporter mouse. VIP:tdTomato-Synaptophysin, SST:tdTomato-Synaptophysin, and PV:tdTomato-Synaptophysin mice were used for the proximity labeling experiments

### Buprenorphine Injections

Two cohorts of 6 C57BL/6 male mice were subcutaneously injected with 0.05 mg/ml buprenorphine or saline control. Mice were kept in their home cages for 4 hours post-injection and then transcardially perfused with 4% paraformaldehyde (PFA) and brains were allowed to postfix for 2 hours in PFA. The cortex was then microdissected and sliced at 60μm before performing proximity labeling as described above with SYT2 as the primary antibody.

### Brain Tissue Sample Preparation

Mice were transcardially perfused with fresh, ice-cold 2% (PFA). Brains were post-fixed in 2% PFA for 2 hours before removal of the cortex through microdissection and vibratome sectioning at 60µm. Slices were collected in 5ml tubes and proximity labeling was performed on free-floating slices.

### Antibody-based Proximity Labeling

Slices were permeabilized with phosphate buffered saline with 0.1% Triton-X (PBS-T) before endogenous peroxidases were quenched with 0.5% hydrogen peroxide (H2O2) for 10 minutes. After washing, slices were blocked for 2 hours in blocking buffer (1% BSA in PBS + 0.1% Tween-20). A primary antibody (rabbit anti-RFP, Rockland 1:500, mouse anti-synaptotagmin 2, Zebrafish International, 1:300, Rabbit PSD-95, 1:2000) was added with an overnight incubation followed by a 4 hour incubation in HRP-conjugated secondary antibody (1:300). Following antibody labeling, samples were incubated for 15 minutes in reaction buffer (100mM borate with 2% dextran sulfate and 0.1% Tween-20) with biotin-tyramide at a ratio of 1:500 (buffer:biotin-tyramide). Biotin-tyramide was synthesized in-house (see *Organic Synthesis of Biotin-Tyramide)*. Biotinylation was initiated through the addition of 0.003% H2O2 and the reaction was allowed to proceed for 1 minute before being quenched with 500mM sodium ascorbate. Samples were rapidly washed in PBST to remove any additional biotin-tyramide before immunohistochemical validation.

### Immunohistochemistry

After biotin labeling, a subset of slices were taken for immunohistochemical validation that biotinylation occurred. Slices were labeled with FITC-streptavidin (Invitrogen, 1:1000), DAPI (1:1000), and an mouse Alexa-568 conjugated secondary antibody (Thermo Fisher, 1:1000). Slices were labeled, mounted, and viewed on a Leica SP8 confocal microscope. The reaction was considered successful if FITC labeling corresponded to the location of the original primary antibody. If the labeling was successful, the remaining free-floating slices were prepared for mass spectrometry.

### Organic Synthesis of Biotin-Tyramide

Biotin-tyramide was synthesized by dissolving 10mg of NHS-Biotin (Thermo Fisher) in dimethylformamide (DMF, Sigma) and allowed to react at room temperature with 10mg tyramine-HCl (Sigma, T90352) dissolved in DMF and triethylamine (Sigma) for 2 hours in the dark. The reaction was stopped through the addition of 100% ethanol. 1 volume of ethyl acetate (Sigma) was then added. Following the reaction, DMF was removed through a liquid-liquid extraction using repeated (3-5) extractions of 5x volume of water, and the biotin-tyramide product was removed and stored at 4°C until use.

### Streptavidin-Enrichment

Slices were homogenized in a 1ml glass Teflon homogenizer. A final concentration of 3% sodium deoxysulfate (Sigma) and 2% sodium deoxycholate (Sigma) were added and samples were heated at 99°^C^ for 1 hour. Samples were centrifuged at 13,000xg for 5 minutes. 150ul of the supernatant was saved for pre-enrichment analysis (“input” LCMS samples), and the remaining supernatant was added to 150µl of Streptavidin M-280 Dynabeads. After incubation at room temperature for 2 hours, the supernatant was removed and saved for analysis (“unbound” LCMS samples). Beads were washed 5x in PBS, and finally resuspended in 300ul of 0.1M ammonium bicarbonate (Sigma).

### On-beads digestion of antibody-based proximity labeled streptavidin pulldowns for analysis by tandem mass spectrometry

Bead-bound samples were reduced in 5 mM dithiothreitol (Thermo Fisher) in 0.1 M ammonium bicarbonate at 23°C for 1.5 h followed by alkylation using 10mM iodoacetamide (Sigma) in 0.1 M ammonium bicarbonate for 45 min at 23°C, covered from light. Samples were proteolyzed with porcine modified sequencing grade trypsin (Promega) at a ratio of 1:20 enzyme:protein at 37°C for 16 hr. Following proteolysis, the peptide-containing supernatant was removed and digestion was quenched with concentrated formic acid to yield a final pH of 3.0. Peptides were desalted on Pierce C18 Peptide Desalting Spin Columns according to manufacturer’s specifications. After desalting, peptides were dried to completion, resuspended in 0.1% formic acid (Pierce) in water, and quantified on a UV-VIS Spectrophotometer using A280/A260 (1 Abs = 1 mg/mL). Injection amounts for mass spectrometry analysis were normalized across all samples based on A280 readings.

### Proteomic Analysis by Ultra-High Performance Liquid Chromatography coupled to Tandem Mass Spectrometry (UPLC-MS/MS)

Peptide samples were subjected to UPLC-MS/MS using a Thermo Scientific Ultimate 3000 RSLCnano ultra-high performance liquid chromatography (UPLC) system coupled to either a Thermo Scientific Q Exactive HF or Orbitrap Eclipse Tribrid mass spectrometer. For Q Exactive HF experiments, desalted peptides resuspended in Solvent A (0.1% formic acid in water) were injected onto a nanoEase M/Z Peptide BEH C18 column (1.7μM, 75μM x 250mm, Waters Corporation) and separated by a 300 nL/min reversed-phase UPLC gradient of 4-30% Solvent B (0.1% formic acid in acetonitrile) over 90 min, 30% to 90% B in 20 min, 90% B hold for 10 min, and return to initial conditions over 2 min with an additional 18 min of column re-equilibration at 4% Solvent B with a total runtime of 150 min. Peptides were ionized directly into the mass spectrometers using positive mode ESI. Q Exactive HF-specific MS parameters included 60,000 resolution, AGC target of 1e6, maximum injection time (MIT) of 60ms, and a mass range of 300-1,800 m/*z*. Data-dependent higher-energy C-trap dissociation (HCD) MS/MS scan acquisition parameters included 15,000 resolution, AGC target of 1e5, MIT of 40 ms, loop count of 15, isolation window of 2.0 m/*z*, dynamic exclusion “on” with a window of 30 sec, normalized collision energy of 27, and charge exclusion “on” for all unassigned, +1, and > +8 charged species. For experiments acquired using the Orbitrap Eclipse Tribrid, either the 150 min UPLC gradient described above was used, or a shortened version with a total runtime of 90 min using identical flow rates and UPLC column was used. Orbitrap Eclipse Tribrid MS parameters were as follows: resolution of 60,000, Standard AGC target, 60 ms MIT, mass range of 300-1800 m/*z*, and 30% RF lens. Data-dependent MS/MS spectra were acquired in the linear ion trap (IT) using collision-induced dissociation (CID) and the following parameters: Dynamic exclusion “on” with a window of 30 sec, charge state inclusion of +2 to +8 charge states only, exclude after n times set to 1, intensity threshold of 5.0e3, cycle time of 3 sec, isolation window of 1.6 m/z, 35% CID collision energy and 10 ms CID activation time, “Rapid” IT scan rate, “Normal” mass range, “Standard” AGC target, and “Dynamic” MIT.

### Peptide identification and label-free quantitation via MaxQuant and data analysis using Scaffold 5

Peptides were identified and quantified for all experiments with the Andromeda search engine embedded in MaxQuant v1.6.0.1 (Cox, J., and Mann, M, 2008). Raw data were searched against the UniProt *Mus Musculus* reference proteome (UP000000589, Accessed 12/12/2017 and updated 05/16/2017) with the fluorescent protein TdTomato sequence appended, plus the MaxQuant contaminants database. Oxidation(M), acetylation(protein N-term), Gln->pyro-Glu, and a Biotin-Phenol(Y) custom modification of 361.1460 Da were set as variable modifications. Carbamidomethylation(C) was set as a fixed modification. Protease specificity was set to trypsin/P with a maximum of 2 missed cleavages. LFQ quantification was enabled and minimum number of amino acids/peptide set to 5. All results were filtered to a 1% false discovery rate at both the peptide and protein levels based on a target/decoy method; all other parameters were left as default values. Peptide/protein identifications by Andromeda and quantification data by MaxQuant were subsequently visualized and analyzed in Scaffold v5.0.1 (Proteome Software) with a minimum of 2 peptides required for identification and average precursor intensity (API) values used for quantification. The mass spectrometry proteomics data have been deposited to the ProteomeXchange Consortium via the PRIDE partner repository with the dataset identifier PXD036293 and reviewer access as follows: **Username:****Password:** 9uS9FYaz”

### Data processing

Scaffold data were imported into MATLAB (version 2021a) for filtering and analysis. Contaminants were removed and final protein list was obtained by removing proteins present in fewer than 3/6 replicates. API values for remaining proteins were averaged across replicates. A background list of proteins known to promiscuously bind to M-280 streptavidin dynabeads was compared to each sample’s protein list and removed from each sample when identified. Protein lists were analyzed using Gene Ontology (Panther, Geneontology.org), Cytoscape, SynGo (SyngoPortal.org), and IPA Ingenuity^tm^. Cytoscape analysis was performed using the ClueGO plugin^35^ using integrated GO terms. Panther Gene Ontology was run using the *Mus musculus* background with a Fisher’s Exact test and Bonferroni correction for multiple hypothesis testing. SynGO results were compared to a background of brain-expressed genes. All SynGO and gene ontology lists were reported in descending order of enrichment and display the top results obtained from the analysis. Network analysis (Fig 1I) was generated using differentially expressed proteins with the greatest fold changes (top 10%, defined by the absolute value of expression ratios) using a force-directed layout^36^ and inverse edge weighting over 1000 iterations.

### Superresolution Microscopy

STED images were collected on an Abberior Instruments Expert Line STED system (Göttingen, Germany) using an Olympus 100x/1.40 UPLANSAPO oil immersion lens. STAR RED was excited with a pulsed 640 nm laser, and STAR ORANGE was excited using a 561 nm pulsed laser. Both dyes were imaged with a pulsed 775 nm depletion laser. STED images were deconvolved to a maximum of 20 iterations and a 0.01 threshold using Huygens software (SVI, Hilversum, Netherlands).

## Notes

### Competing Interest Statement

The authors have declared no competing interest.

